# Neurodevelopmental oscillatory basis of speech processing in noise

**DOI:** 10.1101/2022.01.20.476739

**Authors:** Julie Bertels, Maxime Niesen, Florian Destoky, Tim Coolen, Marc Vander Ghinst, Vincent Wens, Antonin Rovai, Nicola Trotta, Martijn Baart, Nicola Molinaro, Xavier De Tiège, Mathieu Bourguignon

**Affiliations:** Laboratoire de Cartographie fonctionnelle du Cerveau, UNI – ULB Neuroscience Institute, Université libre de Bruxelles (ULB), Brussels, Belgium.; ULBabyLab – Consciousness, Cognition and Computation group, UNI – ULB Neuroscience Institute, Université libre de Bruxelles (ULB), Brussels, Belgium.; Service d’ORL et de chirurgie cervico-faciale, CUB Hôpital Erasme, Université libre de Bruxelles (ULB), Brussels, Belgium.; Department of Radiology, CUB Hôpital Erasme, Université libre de Bruxelles (ULB), Brussels, Belgium.; Department of Functional Neuroimaging, Service of Nuclear Medicine, CUB Hôpital Erasme, Université libre de Bruxelles (ULB), Brussels, Belgium.; BCBL, Basque Center on Cognition, Brain and Language, San Sebastian, Spain.; Department of Cognitive Neuropsychology, Tilburg University, Tilburg, The Netherlands.; Ikerbasque, Basque Foundation for Science, Bilbao, Spain.; Laboratory of Neurophysiology and Movement Biomechanics, UNI – ULB Neuroscience Institute, Université libre de Bruxelles (ULB), Brussels, Belgium.

## Abstract

Humans’ extraordinary ability to understand speech in noise relies on multiple processes that develop with age. Using magnetoencephalography (MEG), we characterize the underlying neuromaturational basis by quantifying how cortical oscillations in 144 participants (aged 5 to 27 years) track phrasal and syllabic structures in connected speech mixed with different types of noise. While the extraction of prosodic cues from clear speech was stable during development, its maintenance in a multi-talker background matured rapidly up to age 9 and was associated with speech comprehension. Furthermore, while the extraction of subtler information provided by syllables matured at age 9, its maintenance in noisy backgrounds progressively matured until adulthood. Altogether, these results highlight distinct behaviorally relevant maturational trajectories for the neuronal signatures of speech perception. In accordance with grain-size proposals, neuromaturational milestones are reached increasingly late for linguistic units of decreasing size, with further delays incurred by noise.

**Teaser:** The neural signature of speech processing in silence and noise features multiple behaviorally relevant developmental milestones

## Introduction

Understanding speech in noise (SiN) is a challenging task, especially for children (*1, 2*). Paradoxically, noise is ubiquitous in children’s lives (e.g., in classrooms, school cafeterias and playgrounds) and has deleterious effects on learning and academic performances (*3*). Still, how the neural mechanisms involved in SiN comprehension mature across development is poorly understood. Characterizing these developmental phenomena appears critical to devise strategies to help children cope with ambient noise in their daily life and to better understand the etiology of learning disorders.

A large body of literature has examined the neurophysiological correlates of SiN processing through investigations of the cortical tracking of speech (CTS) (*4–16*). CTS is the synchronization between human cortical activity and the fluctuations of speech temporal envelope at frequencies that match the hierarchical temporal structure of linguistic units such as phrases/sentences (0.2–1.5 Hz) and syllables/words (2–8 Hz) (*17–25*). Functionally, CTS would subserve the segmentation of these units in connected speech to promote subsequent speech recognition (*18, 19, 24, 26–28*). Importantly, school-age children show reliable CTS (*21, 29, 30*) that is however lower at the syllabic level compared to adults (*5*). In SiN conditions, children’s and adults’ cortical activity preferentially tracks the attended speech rather than the global sound (*4, 6, 10, 31*), suggesting that CTS is modulated by endogenous attentional components and plays a role in segregating the attended linguistic signal (*5–8, 10, 12, 13*). However, the fidelity of the tracking decreases with increasing noise intensity in adults and more so in children (*4–6*), especially when the noise is concurrent speech babble (*32*) as opposed to non-speech noise such as white or spectrally-matched noise. Of note, the visual speech signal (comprising the articulatory movements of a talker) boosts CTS in adults (*33–35*), especially in noise conditions (*31, 36, 37*), and in children, at least in babble noise conditions (*32*). Importantly, the modulations of CTS we have just outlined mirror tangible behavioral effects. That is, SiN perception and comprehension *(i)* decline with increasing noise level (*4–6*), *(ii)* are more affected by babble than non-speech noise in adults and especially in children (*38–40*), *(iii)* improve until late childhood, if not until adolescence in babble noise conditions (2,4,5), and *(iv)* benefit from visual speech (*41, 42*) since infancy (*43, 44*), but increasingly more as age increases (*45, 46*).

Overall, these data suggest (i) that different aspects of CTS, whose behavioral relevance is well demonstrated, undergo different developmental trajectories, and (ii) that these trajectories depend on noise properties and availability of visual speech information. However, since previous studies focused on restricted age ranges and noise conditions, a detailed characterization of these trajectories is still lacking. The present magnetoencephalography (MEG) study aims at filling this gap by outlining the developmental trajectory of phrasal and syllabic CTS and speech comprehension, from early school age to early adulthood in various noise conditions, with or without visual speech information. Our research hypotheses were guided by *grain-size* proposals according to which children develop awareness of increasingly smaller phonological units with age (*47, 48*). Extrapolating these proposals to supra-phonological units (*19, 49*), we hypothesized that the cortical tracking of large units such as phrases and sentences would mature faster during development compared with the tracking of smaller units such as syllables. Also, since coping with noise and leveraging visual speech information require the development and integration of additional mechanisms subtended by high-order associative neocortical areas that mature during late childhood (*50*), we hypothesize that corresponding developmental trajectories would be further delayed.

## Results

We recorded brain activity with MEG in 144 participants (77 females) aged 5.3–27.0 years while they were attending to 4 videos lasting ∼6 min each, following the same experimental procedure as in a previous study by our group (*32*). Videos consisted of audiovisual recordings of native French-speaking narrators telling children’s fairy tales. Fig. 1 illustrates the time-course of a video stimulus. Each video featured 9 conditions: 1 noiseless and 8 SiN with 3 dB signal-to-noise ratio (SNR; for the motivation behind this SNR selection, see *Stimuli* subsection in *Materials and Methods*) resulting from the combination of 4 types of noises (least-energetic non-speech, most-energetic non-speech, different-gender babble and same-gender babble) with 2 visual conditions (with or without visual speech inputs). The different- and same-gender babble noises introduced informational interferences and a similar degree of energetic masking (*32*). Their distinction is however relevant since speech intelligibility is generally better when attended and interfering speech are uttered by different-gender talkers compared to same-gender talkers (*51, 52*), because on average, voice fundamental frequency and vocal tract length differ between males and females (*52, 53*). The least- and most-energetic non-speech noises introduced a degree of energetic masking in accordance with their naming but no informational interference. Forty yes/no questions (10 per video) assessed participants’ comprehension of the stories in each condition.

**Fig. 1.**
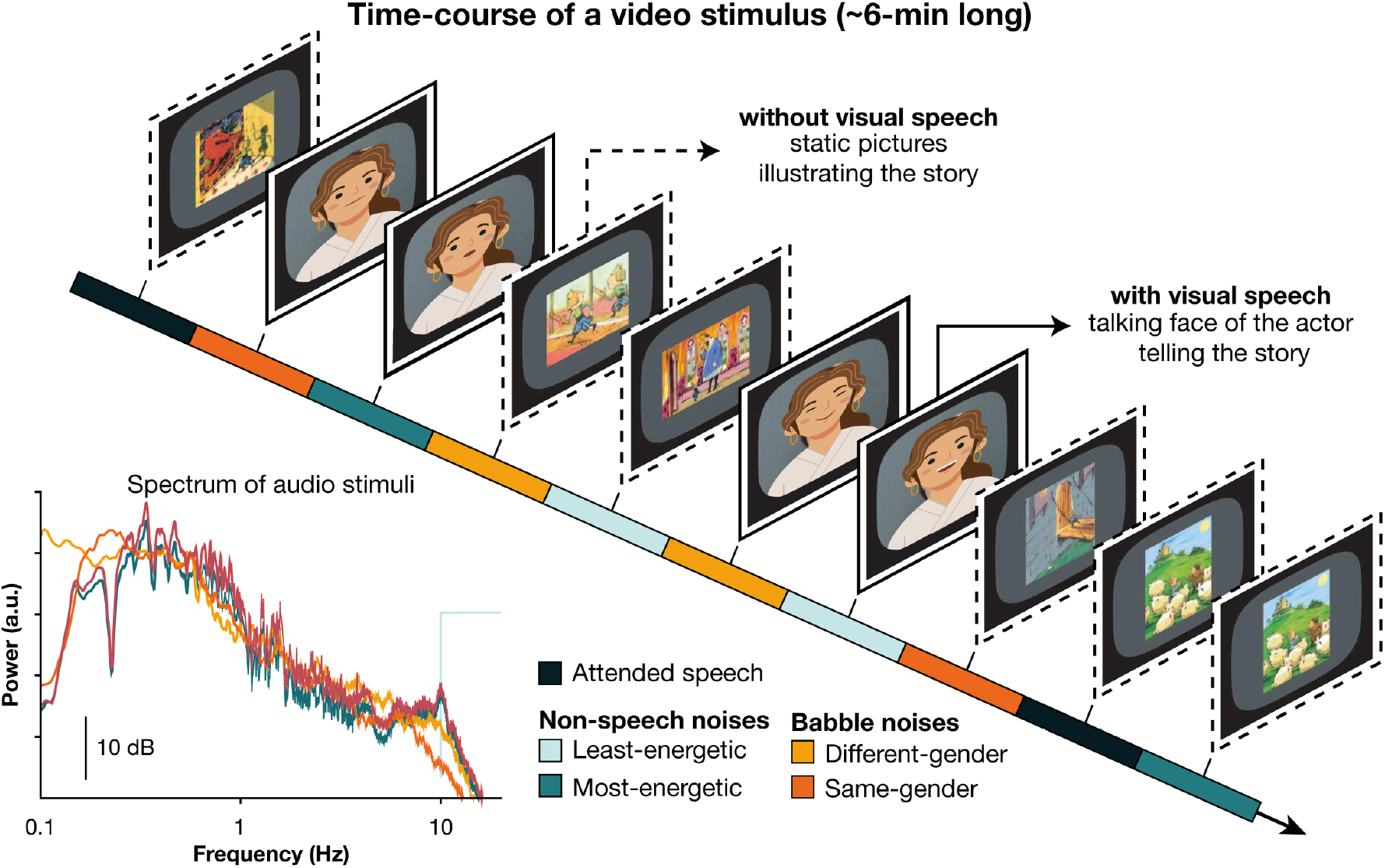
Illustration of the time-course of a video stimulus. Videos lasted approximately 6 min and were divided into 10 blocks to which experimental conditions were assigned. There were two blocks of the noiseless condition, and eight blocks of speech-in-noise conditions: one block for each possible combination of the four types of noise and the two types of visual display.

For some of the upcoming analyses, participants were arranged in 5 age groups of roughly equal size (5–7 years, *n* = 31; 7–8.5 years, *n* = 34; 8.5–11.5 years, *n* = 28; 11.5–18 years, *n* = 26; and 18–27 years, *n* = 25). Note that most of our participants were aged below 12, since SiN capacities essentially develop before that age. As a consequence, the 3 age groups of school-age children (5–7 years, 7–8.5 years and 8.5–11.5 years) span a narrower age range than the groups of teenagers (11.5–18 years) and young adults (18–27 years).

### How does the cortical tracking of speech evolve with age in the absence of noise?

For each condition, we regressed the temporal envelope of the attended speech on MEG signals with several time lags using ridge regression and cross validation (see Methods for details) (*54*). The ensuing regression model was used to reconstruct speech temporal envelope from the recorded MEG signal. Such analysis is known as reconstruction accuracy (*54*). CTS was computed as the correlation between the genuine and reconstructed speech temporal envelopes. We did this for MEG and speech temporal envelope signals filtered at 0.2–1.5 Hz (phrasal rate, which also englobes sentential rate) (*4, 32, 55*) and 2–8 Hz (syllabic rate, which also englobes word rate) (*10, 14, 31, 56*) and for MEG sensor signals in the left and right hemispheres separately. We assessed the left- and right hemispheres separately because CTS is hemispherically asymmetric both in noiseless and SiN conditions (*6*). We first evaluated with an ANOVA if CTS in the noiseless condition depended on the hemisphere and on the age group. A summary of the results for phrasal and syllabic CTS are presented in Supplementary material (Table S1). Phrasal CTS was higher in the right (0.44 ± 0.09; mean ± SD across subjects) than in the left hemisphere (0.40 ± 0.08), and was not modulated by age. Syllabic CTS was also higher in the right (0.092 ± 0.036) than in the left hemisphere (0.079 ± 0.034), but a significant interaction with age indicated that left- and right-hemisphere CTS underwent different developmental trajectories.

We next used Spearman correlations and a model-fitting approach to better understand how age impacted syllabic CTS. Fig. 2 and Table 1 illustrate the developmental trajectory of syllabic CTS. While both left- and right-hemisphere CTS were similar in children aged below 7, a maturation process starting at 7.7±1.7 years increased right-but not left-hemisphere CTS by ∼30 %, plateauing at 10.4±1.9 years.

**Figure 2.**
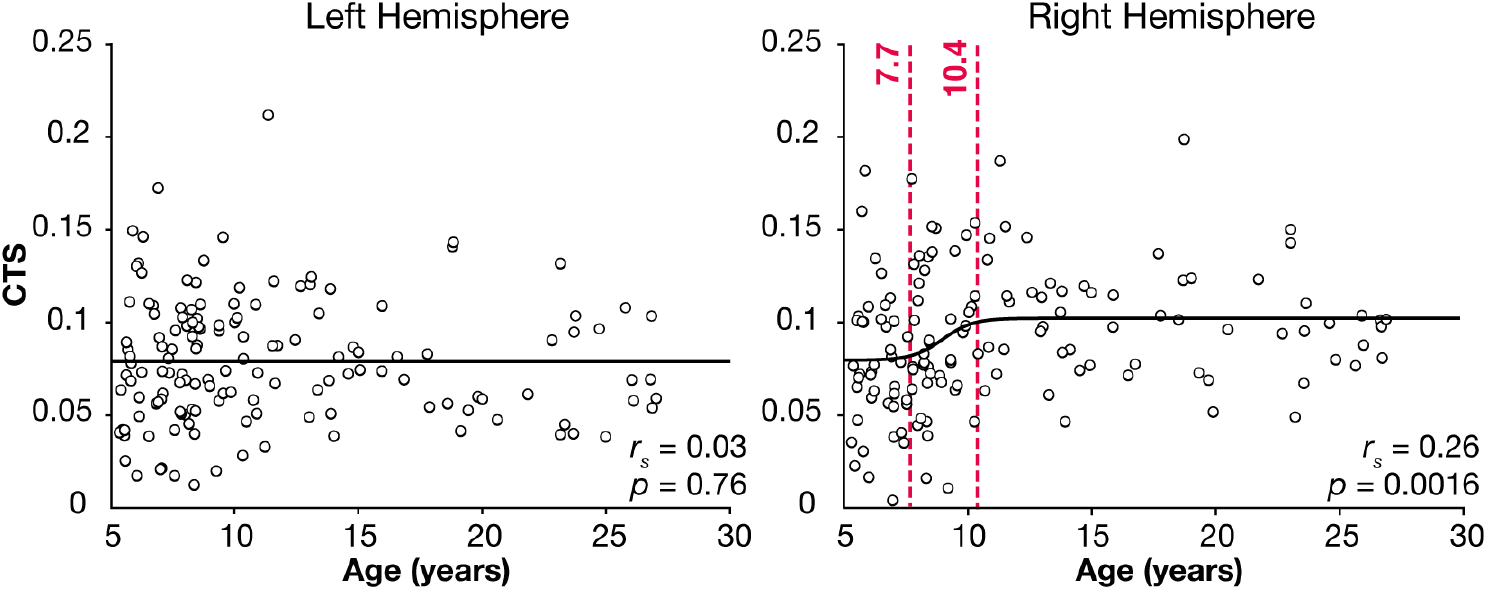
Dependence on age of syllabic CTS in the noiseless condition. Dashed red lines indicate the beginning and the end of the maturation process.

**Table 1.** Parametric models of the dependence on age of (n)CTS values and speech comprehension scores. The number of participants (n) on which models were fitted was 144 for (n)CTS values and 142 for speech comprehension scores. Values of normalized CTS (nCTS) were pooled across conditions with and without visual speech, and across least- and most-energetic conditions. Values of phrasal nCTS were further pooled across hemispheres.

|  | Constant vs. Linear |  | Linear vs. Logistic |  | Constant vs. Logistic |  | Model |
| --- | --- | --- | --- | --- | --- | --- | --- |
|  | *F*(1,*n*-2) | *p* | *F*(2,*n*-4) | *p* | *F*(3,*n*-4) | *p* |  |
| Syllabic CTS in noiseless |  |  |  |  |  |  |  |
| left hemisphere | 0.17 | 0.69 | 1.87 | 0.16 | 1.30 | 0.28 | 0.080 |
| right hemisphere | **3.98** | **0.048** | **4.18** | **0.017** | **4.17** | **0.0073** | $0.079 + 0.023 / (1 + \exp(-1.6(\text{age} - 9.0)))$ |
| Phrasal nCTS |  |  |  |  |  |  |  |
| non-speech | 2.06 | 0.15 | 2.09 | 0.10 | 2.09 | 0.13 | $-0.01 + 0.02 / (1 + \exp(-787(\text{age} - 8.6)))$ |
| babble | **31.4** | **<0.0001** | **20.5** | **<0.0001** | **12.5** | **<0.0001** | $-0.37 + 0.27 / (1 + \exp(-1.1(\text{age} - 7.3)))$ |
| Syllabic nCTS |  |  |  |  |  |  |  |
| non-speech, left hem | **4.94** | **0.028** | 0.77 | 0.47 | **2.15** | **0.096** | $-0.035 + 0.0039 \times \text{age}$ |
| babble, left hem | **7.99** | **0.0054** | 1.51 | 0.22 | **3.69** | **0.014** | $-0.25 + 0.0067 \times \text{age}$ |
| non-speech, right hem | 1.86 | 0.17 | 0.24 | 0.79 | 0.77 | 0.51 | 0.027 |
| babble, right hem | **4.18** | **0.043** | 0.002 | 1.00 | 1.38 | 0.25 | $-0.20 + 0.0049 \times \text{age}$ |
| Speech (story) comprehension scores |  |  |  |  |  |  |  |
| noiseless | **20.7** | **<0.0001** | **18.6** | **<0.0001** | **21.0** | **<0.0001** | $0.79 + 0.16 / (1 + \exp(-770(\text{age} - 6.9)))$ |
| noise | **35.0** | **<0.0001** | **94.3** | **<0.0001** | **76.3** | **<0.0001** | $-28755.83 + 28756.78 / (1 + \exp(-0.87(\text{age} + 7.8)))$ |
| difference Lip-Vid | **3.07** | **0.082** | **2.84** | **0.062** | **2.94** | **0.035** | $-0.065 + 0.086 / (1 + \exp(-1.2(\text{age} - 7.3)))$ |

### How does noise impact the cortical tracking of speech, and how does this impact evolve with age?

We first evaluated with an ANOVA whether phrasal and syllabic normalized CTS (nCTS) in noise conditions depend on noise properties, hemisphere, visibility of the talker’s lips and whether they evolve with age. The nCTS is a contrast between CTS in SiN and noiseless conditions (see Methods) that takes values between –1 and 1, with negative values indicating that the noise reduces CTS (*32*). Such contrast presents the advantage of being specific to SiN processing abilities by factoring out the global level of CTS in the noiseless condition.

Fig. 3 summarizes the results for phrasal and syllabic nCTS (see also Table S2). Overall, the impact of the different types of background noises was similar for phrasal and syllabic nCTS: while non-speech noises did not affect much CTS (nCTS was close to zero), babble noises substantially reduced CTS compared to non-speech and noiseless conditions. Contrastingly, the level of energetic masking introduced by either type of noise only mildly affected the nCTS. Such pattern was observed for both hemispheres, and irrespective of the availability of visual speech information. Nevertheless, phrasal nCTS in babble noise conditions was higher in the left than right hemisphere (−0.16 ± 0.20 vs. −0.20 ± 0.19) while the reverse was true for syllabic nCTS in all noise conditions (−0.10 ± 0.24 vs. −0.07 ± 0.22).

**Fig. 3.**
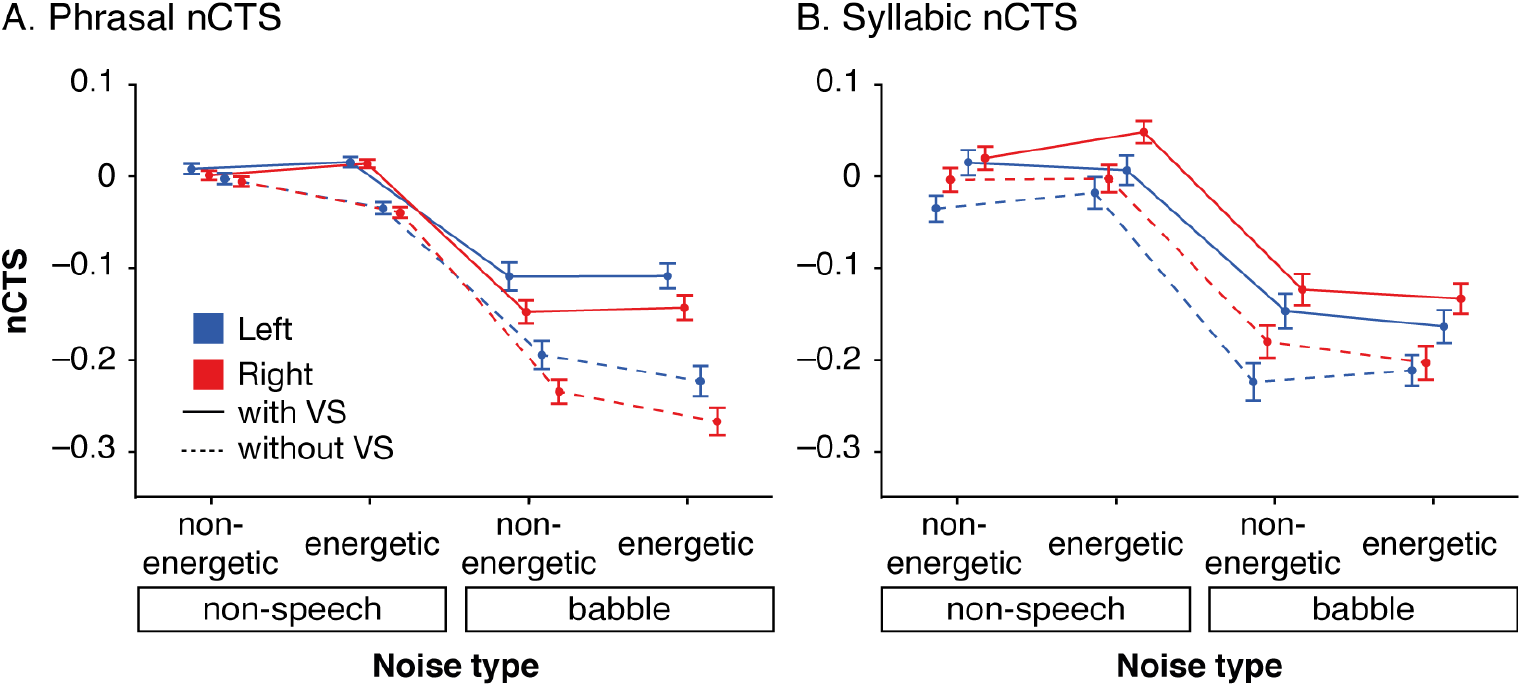
Impact of the main effects on nCTS at phrasal (A) and syllabic rates (B). Mean and SEM values are displayed as a function of noise properties. The four traces correspond to conditions with (connected traces) and without (dashed traces) visual speech (VS), within the left (blue traces) and right (red traces) hemispheres. nCTS values are bounded between –1 and 1, with values below 0 indicating lower CTS in speech-in-noise conditions than in noiseless conditions.

A beneficial effect of visual speech information was observed in all noise conditions except in the least challenging one (i.e., least-energetic non-speech) for phrasal nCTS, and in all noise conditions for syllabic nCTS. The way visual speech information modulated nCTS was stable across the age range (*ps* > 0.2 for interactions involving age and type of visual input).

Critically, the way different noises impacted both phrasal and syllabic nCTS differed between age groups (i.e., significant interactions involving age and noise). To better understand how nCTS evolved with age, we relied on the same approach applied to CTS in the noiseless condition, which involves Spearman correlations and model fitting.

Fig. 4 and Table 1 present the results for nCTS pooled across least- and most-energetic noises. The detailed results for all noise conditions separately are presented in Supplementary material (Fig. S1, and Table S3).

**Fig. 4.**
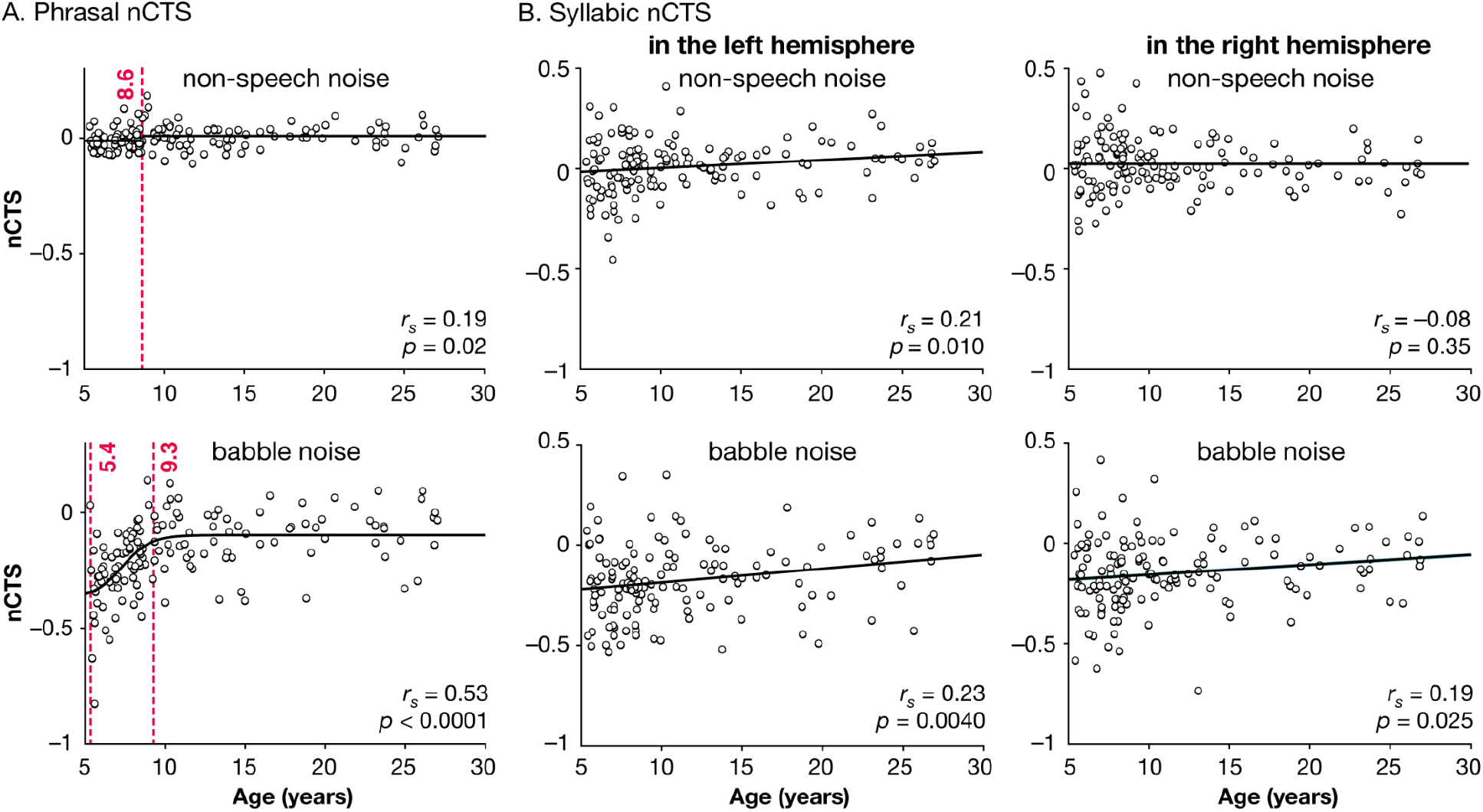
Dependence on age of phrasal (A) and syllabic (B) nCTS. nCTS was pooled across conditions with and without visual speech, and across least- and most-energetic conditions. Phrasal nCTS was further pooled across hemispheres. Dashed red lines indicate the beginning and end of the maturation process.

Phrasal nCTS increased with age for both non-speech and babble noises. The modulation in CTS was however only marginal in non-speech noise conditions (4.2% with a transition at 8.6 ± 0.0 years following a logistic model), to the point that our model-fitting approach did not deem the age-dependent models better than a constant model. This suggests a minimal evolution of phrasal nCTS in non-speech noises, at least at a SNR of 3 dB. As a slight nuance, the evolution was clearer when considering the most-energetic non-speech noise, and fully absent for the least-energetic non-speech noise (Fig. S1 and Table S3). In contrast, a clear maturation process starting at 5.4 ± 1.6 years increased phrasal nCTS in babble noises by ∼79 %, with a plateau at age 9.3 ± 0.9 years.

Syllabic nCTS also increased with age, following linear trajectories, with different patterns observed for non-speech and babble noises in the left and right hemispheres. That is, syllabic nCTS increased with age in both hemispheres for babble noise, and only in the left hemisphere for non-speech noise.

### How does speech comprehension evolve with age, in noiseless and noise conditions?

Fig. 5 illustrates speech comprehension abilities in the different conditions, which were assessed using yes/no forced-choice questions after each video. Comprehension scores were computed as the percentage of correct answers to 4 questions in each noise condition (or 8 in the noiseless condition).

**Fig. 5.**
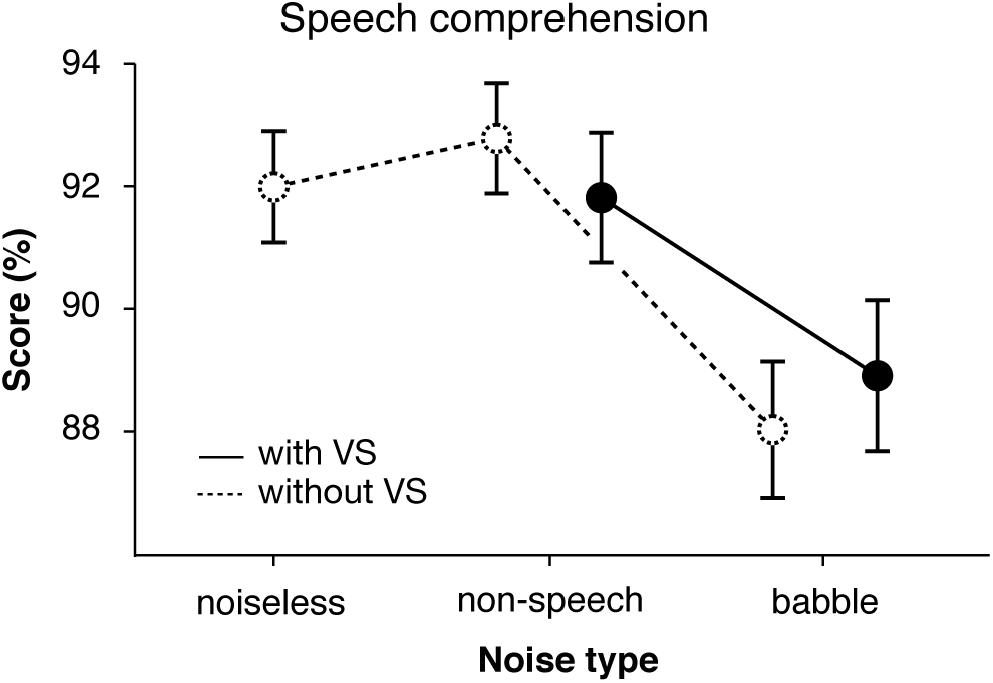
Impact of the noise condition and of the presence or absence of concomitant visual speech on the speech comprehension scores, pooled across age groups. Vertical bars indicate SEM values.

Comprehension of the stories differed between age groups in noiseless (*F*(4, 137) = 14.0, *p* < 0.0001) and noise conditions (*F*(4,137) = 26.1, *p* < 0.0001), improving with age in both cases. The model fitting approach identified a sharp transition at 6.9 years for comprehension in silence, and absurd values (negative transition age) for comprehension in noise. Comprehension in noise conditions was also affected by noise properties (*F*(3,411) = 7.31, *p* < 0.0001). It was better when non-speech (92.3 ± 15.3 %) compared to babble noises (88.5 ± 17.8 %) were presented in the background. In fact, comprehension in non-speech noise conditions was similar (*t*(141) = −0.37, *p* = 0.72) to that in the noiseless condition (92.0 ± 11.2 %), indicating that non-speech noise had no detrimental effect on the comprehension of the story (for our 3-dB SNR level). Finally, a marginally significant interaction between visual input and age group (*F*(4,137) = 2.34, *p* = 0.059) suggested that the benefit of visual speech for speech comprehension differed between age groups. The exploration of the boost in comprehension induced by visual speech is presented in Supplementary Fig. S2. No other significant effects were disclosed (*p* > 0.1).

### Behavioral relevance of the cortical tracking of speech

We next appraise the behavioral relevance of the tracking measures showing significant maturation effects (i.e., syllabic CTS in the right hemisphere, phrasal nCTS in babble noise, and syllabic nCTS in each hemisphere and in non-speech and babble noise conditions separately). For this, we correlated (n)CTS measures and comprehension scores, after removing the—potentially non-linear—effect of age.

In the noiseless condition, this analysis revealed no significant association between syllabic CTS and speech comprehension (*ps* > 0.3 for left- and right-hemisphere CTS).

In SiN conditions, this analysis revealed a positive correlation between phrasal nCTS (averaged across hemispheres) and speech comprehension in babble noise conditions (*r*_S_ = 0.22; *p* = 0.0074; see Fig. 6), and no significant association between syllabic nCTS and speech comprehension, neither in non-speech (*ps* > 0.5 for left- and right-hemisphere CTS) nor in babble noise conditions (*ps* > 0.9).

**Fig. 6.**
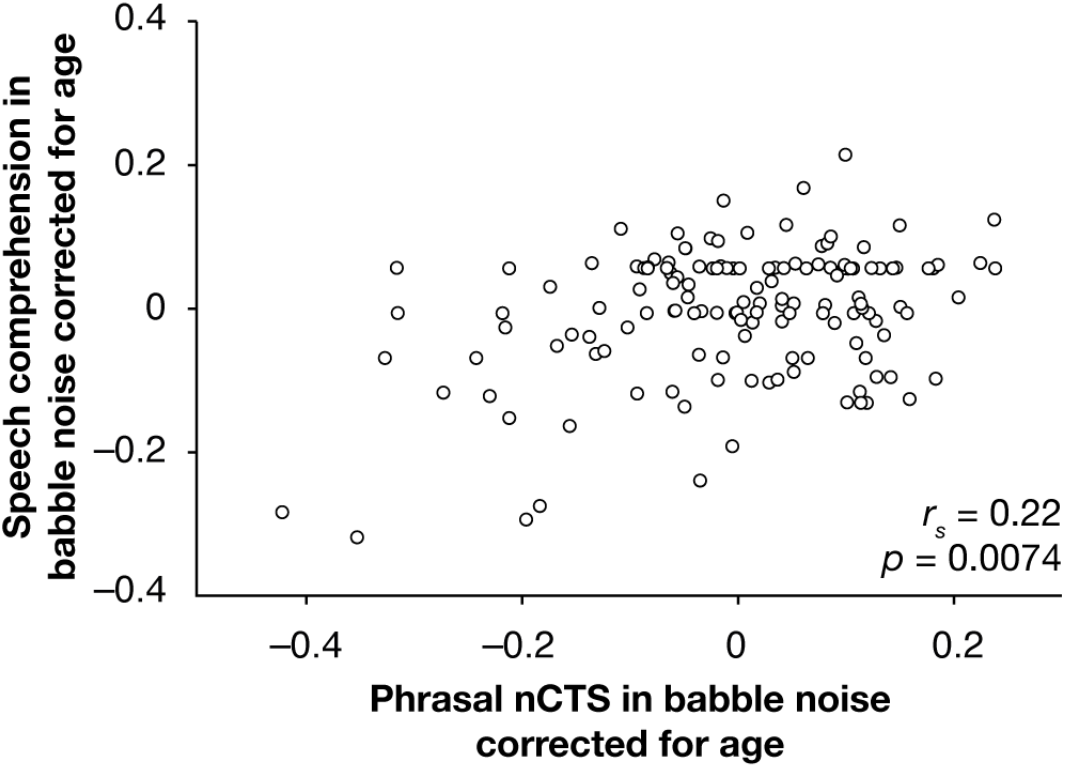
Behavioral relevance of phrasal nCTS in babble noise.

### Sources of the cortical tracking of speech

We next identified the cortical sources underlying the CTS. Source activity was reconstructed with minimum norm estimate, and CTS was assessed for each source separately with a measure of reconstruction accuracy akin to that used in the previous sections (*54*). For each combination of age group (below 7, 7–8.5, 8.5–11.5, 11.5–18, and 18-27), condition, and frequency range (phrasal and syllabic), we retained the coordinates of significant local maxima of CTS. Since local maxima indicate the presence of underlying sources (and colocalize with them), we will term them the sources of CTS.

Fig. 7 presents the grand average CTS map (mean across all factors), together with the location of the significant sources of CTS in all conditions. Globally, sources of phrasal CTS localized bilaterally in the mid-superior temporal gyrus (STG), in the ventral part of the inferior frontal gyrus (IFG; in partes opercularis, triangularis and orbitalis) and precentral gyrus and, to a lower extent, in posterior temporal regions. Sources of syllabic CTS essentially localized bilaterally in tight clusters centered around Heschl gyrus and in the anterior part of the IFG (partes orbitalis and triangularis) and, for few of them, in the temporoparietal junction (TPJ) and inferior part of the precentral gyrus.

**Fig. 7.**
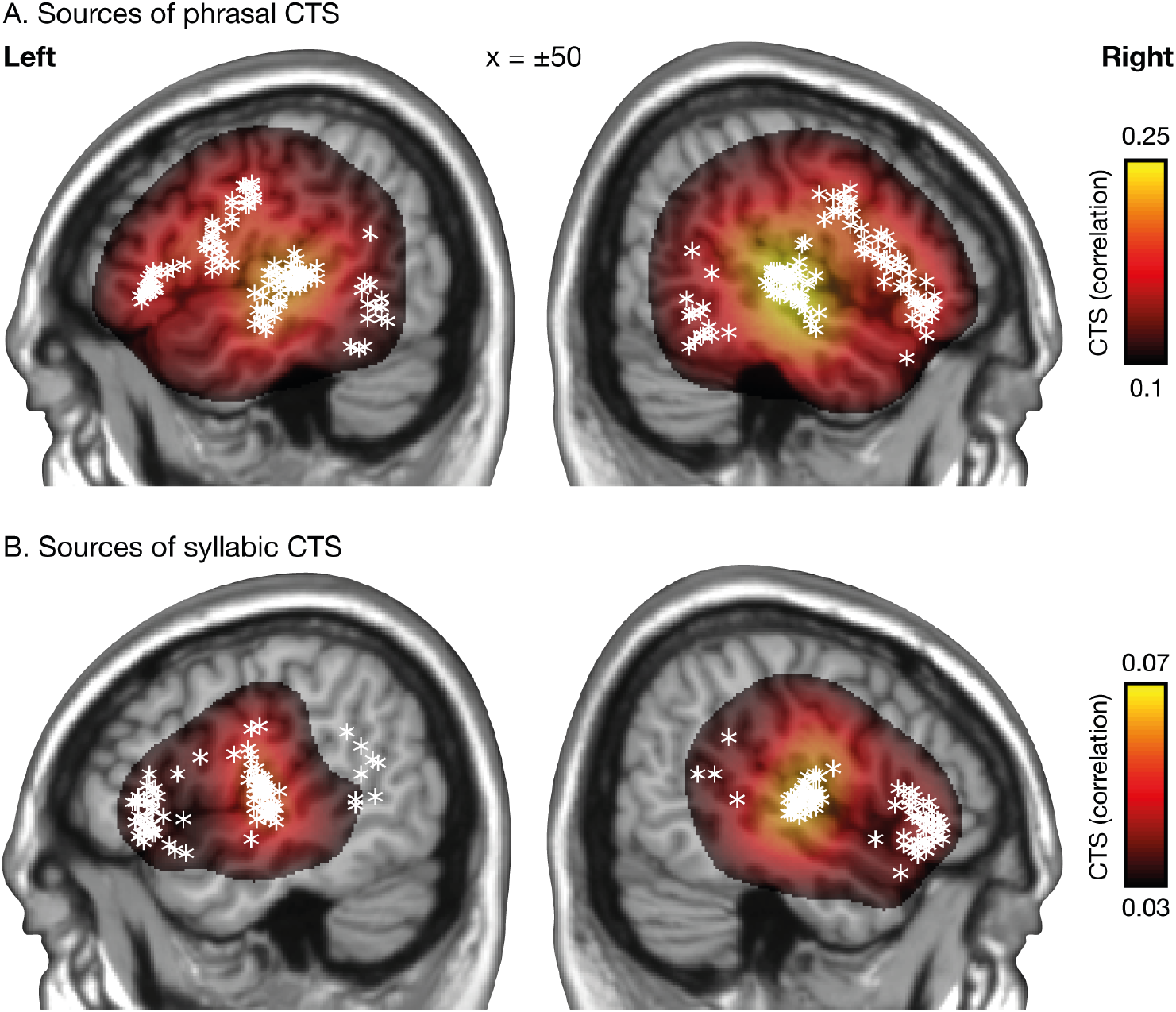
Sources of phrasal (A) and syllabic (B) CTS in the left and right hemispheres. The overlays present the mean CTS values across all conditions and participants (regardless of age). Values at MNI coordinates |X| > 25 mm were projected orthogonally onto the parasagittal slice of coordinates |X| = 50 mm. The location of each significant source of CTS in each condition and age group is indicated with a white star (with the same projection scheme).

Next, we evaluated for each frequency range if sources of CTS tended to cluster according to age group, or different noise properties.

First, sources of phrasal CTS had among their 10 closest neighbors 62.9 % more sources for the same age group than expected by chance (*p* < 0.0001). To better understand this effect, Fig. 8A presents the sources of phrasal CTS color-coded by age group. Sources in the right hemisphere tended to localize more posteriorly with increasing age. Other differences were more subtle and not characterized by clear age gradients or source presence from or before a given age (e.g., sources in the right posterior temporal region were not seen in age groups of 7–8.5 years and 18–27 years; sources in precentral gyri were not seen in age groups of 5-7 years and 11.5–18 years).

**Fig. 8.**
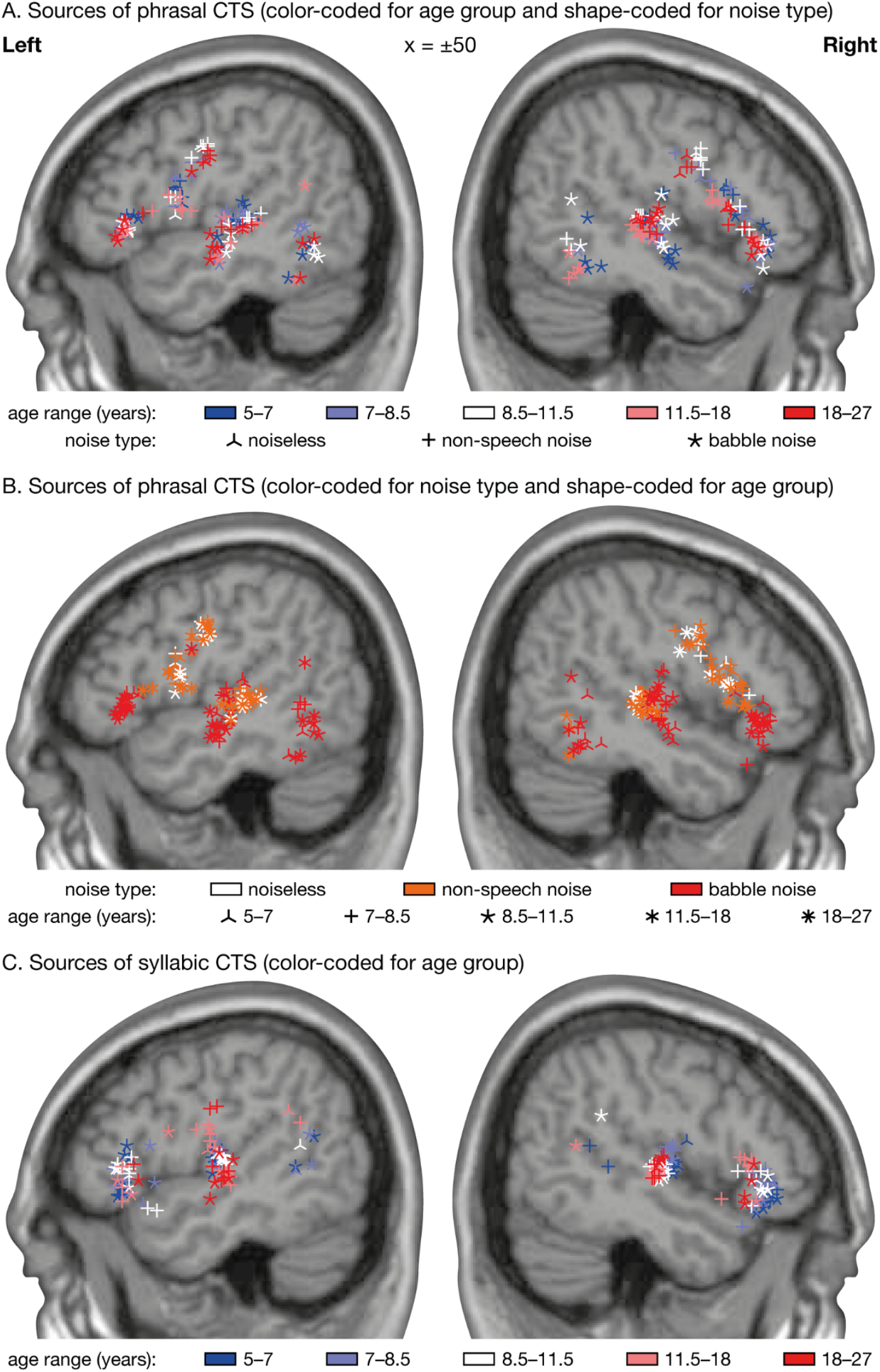
Sources of CTS color-coded for age group (A, phrasal; C, syllabic) and for the informational property of the noise (B, phrasal); the other property being shape-coded.

Second, sources of phrasal CTS for (i) babble noise conditions on the one hand, and (ii) non-speech noise and noiseless conditions on the other hand, had among their 10 closest neighbors 66.1 % more sources for the same category (i.e., i or ii) than expected by chance (*p* < 0.0001). Fig. 8B presents the sources of phrasal CTS color-coded for the informational property of the noise. Sources in bilateral STG and IFG were more anterior for babble noise conditions than for non-speech noise and noiseless conditions. Sources in bilateral IFG localized in the pars orbitalis/triangularis for babble noise conditions, and in the pars triangularis/opercularis as well as in the inferior part of the precentral gyrus for non-speech noise and noiseless conditions. Finally, all sources in the posterior temporal areas were from babble noise conditions except for 3 right-sided sources from non-speech noise conditions.

Third, sources of syllabic CTS had among their 10 closest neighbors 76.8 % more sources for the same age group than expected by chance (*p* < 0.0001). To better understand this effect, Fig. 8C presents the sources of syllabic CTS color-coded by age group. Paralleling the effect found for phrasal CTS, sources of syllabic CTS in the right hemisphere tended to localize increasingly more posteriorly with increasing age. Other subtler effects included the absence of source in TPJ in the oldest age group (18–27), and more scattered source distributions along the ventrodorsal axis in the left Heschl gyrus in the two oldest age groups (11.5–18 and 18–27; sources reached the ventral bank of the STG and the ventral part of the postcentral gyrus).

## Discussion

This study characterizes the maturation of neurophysiological markers of the perception and understanding of natural connected speech in silence and in noise with or without visual speech information. Our results highlight that while phrasal CTS in quiet conditions is adult-like from at least 5 years of age, syllabic CTS matures later in childhood. We also demonstrated two distinct neuromaturational effects related to the ability to perceive speech in babble noise: while the ability to maintain phrasal CTS matures rapidly between ∼5 and ∼9 years, a much slower maturation process improves the ability to maintain syllabic CTS in babble noise through childhood and into early adulthood. Visual speech information increased phrasal CTS mainly in babble noise conditions and syllabic CTS similarly in all noise conditions. These effects were not modulated by age. The results also reveal a limited impact of age on the cortical sources of phrasal and syllabic CTS.

### Increase of syllabic but not phrasal CTS in silence during childhood

Our data revealed different developmental trajectories related to the capacity of the brain to track the fluctuations of speech temporal envelope at different frequencies, in noiseless conditions. While phrasal CTS is adult-like from at least 5 years of age, syllabic CTS matures later, between 7.5 and 10.5 years. This difference in developmental trajectory is well in line with grain-size proposals (*47, 48*) extended to linguistic units we have proposed. Indeed, phrasal CTS is considered to partly reflect prosodic (*17, 57*) and linguistic (*27, 58*) processing of large speech units. Contrastingly, syllabic CTS would reflect parsing of syllable rhythms (*27*), and the sensitivity to this basic unit of speech (*59*) would be at the basis of efficient phonemic processing (*60*). Importantly, syllabic CTS is considered a lower-level process tightly related to the acoustic features of the auditory input (*28, 61*).

Our finding that phrasal CTS is adult-like from at least 5 years of age supports the view that tracking of slow phrasal and prosodic stress patterns is a foundational process that might be present since birth (*62*), and remains stable across middle adulthood (*63*). This result is not so surprising if one embraces the view that phrasal CTS partly underpins prosodic speech processing (*17*). Indeed, young infants already use such information to parse speech into words and phrases (*64*). Accordingly, other neurophysiological markers of brain processing of prosody in speech (i.e., closure positive shift) were reported in 6-months old in relation to brief pause detection but also to pitch variations (*65*). The result of stable phrasal CTS from 5 years on is also compatible with the view that phrasal CTS reflects lexical and syntactic processing. Indeed, typically developing children of that age possess basic syntactic skills (*66*) such as the ability to understand relative clauses (*67, 68*).

We found evidence for a developmental boost in syllabic CTS in quiet conditions in the right-(but not left-) hemisphere. This boost mostly occurred between the age of 7.5 and 10.5 years, and signified the start of right-hemisphere dominance for syllabic CTS. This transition suggests that, although operational early in life (*62, 69*), temporal parsing of the speech signal at the syllabic level refines with brain maturation. And indeed, children aged below 10 are less accurate than adults at identifying syllable boundaries when these are defined only by amplitude modulations in speech temporal envelope (*70*). The right-hemispheric dominance in noiseless conditions observed for syllabic CTS after age ∼10 and for phrasal CTS is consistent with previous findings in children and adults (*5, 17, 19, 21, 30, 61, 71, 72*). It is even at the core of the *asymmetric sampling in time hypothesis*, which argues that prosodic and syllabic information are preferentially processed in the right hemisphere, while phonemic information is preferentially processed in the left hemisphere or bilaterally (*73*). As previously argued (*74*), the fact that language brain functions become asymmetric in the course of development suggests asymmetry is a hallmark of maturity.

### Development of the neurophysiological basis of speech perception in noise

Our results highlight two distinct neuromaturational effects related to the ability to perceive speech in babble noise. First, the ability to maintain phrasal CTS in babble noise matures rapidly between ∼5 and ∼9 years, with a marked transition at age ∼7. Second, a much slower maturation process—best characterized by a linear progression with age—improves the ability to maintain syllabic CTS in babble noise through childhood and into early adulthood. Following the rationale developed in the previous subsection, our results indicate a rapid maturation at age ∼7 of the neurophysiological mechanisms at play in processing prosodic and suprasegmental linguistic information in natural connected speech in babble noise, and a slower, progressive maturation into early adulthood of the mechanisms involved in the extraction of hierarchically lower syllabic, phonemic or even acoustic information from speech in babble noise. This is well in line with our working hypotheses: neuronal processing of larger linguistic units (words and phrases) develops before that of smaller syllabic units, and coping with noise necessitates additional processes that mature later on.

The maturational time-course of the ability to maintain phrasal CTS in babble noise closely parallels that of the ability to recognize words in the presence of two-talker speech. The latter improves progressively from 5 to 10 years of age, reaching adult-like levels at age 11 (*40, 75, 76*). This maturation trajectory is specific to informational noise since speech recognition in speech-shaped noise is close to adult-like already at age 5 (*40, 75*), as was phrasal CTS in non-speech noise in our data. This suggests that maintenance of phrasal CTS reflects a range of processes involved in the ability to perceive and understand linguistic chunks larger than syllables or words in babble noise. This interpretation is further supported by the similar developmental trajectory of our measures of SiN comprehension, and by the finding that CTS resistance to babble noise is positively related to speech comprehension after having accounted for age.

The degree of maintenance of phrasal CTS in babble noise could actually underpin the maturation of auditory stream formation, which is the process of grouping together sounds from the same source (*77*). Forming auditory streams is a challenging aspect of speech perception in noise, and failure in forming streams seems to explain the behavioral difficulties understanding speech from among two same-gender talkers (*52*). From the point of view of development, the ability to form auditory streams based on frequency separation appears to be immature at age 5-8, and adult-like at age 9–11 (*78*). Since these developmental milestones match well with those found for phrasal CTS in babble noise in our study, the way babble noise impacts phrasal CTS could represent an electrophysiological signature of the ability to form auditory streams.

The slow maturation of the ability to maintain syllabic CTS in noise closely parallels the evolution of phonemic perception in noise. Although such slow evolution was not evident in our phonemic perception test (see Supplementary Material), it is clearly seen in normative data for this test where twice more items were used to assess an even larger sample of participants than ours (*79*). In that study, phonemic perception in noise improved steadily from age 5, topped in the 15-19 year group, and then decreased in the subsequent age ranges, the first of which was overly broad (20-49 years) unfortunately. Our data therefore provide a neurophysiological ground for the slow maturation of phonemic perception in noise. It also suggests a more important role of the left hemisphere since maturation was not observed in the non-speech noise condition in the right hemisphere. This is well in line with the classical dominant role of the left hemisphere for language comprehension.

### Impact of visual speech on CTS

Our data did not reveal any evidence for a maturation of the boost in CTS afforded by visual speech across the tested age range. This is somewhat surprising since audiovisual integration processes mature rather slowly. For some tasks, adult-like performance is reached after age 12 (*45, 46, 80*).

The analysis of our comprehension scores hinted at a transition between age 6 and 9 in the ability to leverage visual speech to enhance comprehension. This is in-line with the observation that at around 6.5 years of age, children start to benefit from having phonetic knowledge about severely degraded speech sounds when asked to match such a sound with a visual speech video (*81*). Possibly then, a CTS boost induced by visual speech may be driven by audiovisual congruence detection, an ability that is already observed at 2 months of age (*43, 44*). Although this suggestion provides an interesting avenue for future work, it is currently rather tentative as processing congruence in audiovisual speech seems to start at around 200 ms (*82*), can take several hundreds of milliseconds (*81, 83*) and therefore overlaps in time with other processes (such as processing of lexico-semantic information) that are difficult to disentangle.

In the S1 discussion, we elaborate further on the beneficial effects of visual speech on CTS we observed across all age ranges.

### Recruited neural network and impact of maturation and noise

Our results showed that source configuration was affected by age for both phrasal and syllabic CTS, and by informational noise properties for phrasal CTS.

The effect of age for both phrasal and syllabic CTS appeared to be mainly explained by an anterior-to-posterior shift (of about 1 cm) of right-hemisphere sources from youngest (5–7 years) to oldest (18–27 years) age groups. Whether this shift reflects a genuine developmental effect is difficult to tell since changes in brain anatomy from childhood to adulthood induce small, but consistent, age-dependent errors in the normalization of individual brains to a template (*84*). Besides these unclear effects of age, our results rather emphasize the close similarity in location of cortical generators of CTS across the investigated age range. This is in line with a host of findings indicating that the architecture of the language network is settled from age 3, with subsequent maturation essentially refining bottom-up communication and specialization of each node of the network (*85*).

In S2 Discussion, we discuss the interesting effect of noise properties on the configuration of CTS sources.

### Limitations

We manipulated several properties of the noise but not all of those known to impact SiN perception. It is therefore worth noting that the developmental trajectories we report for CTS resistance to noise are valid only for the conditions we explored, and might be affected by other aspects of the listening condition, much like the maturation timeline of behavioral effects depends on the number of speakers making up the background noise (*86*), noise intensity (*87*), or availability of spatial cues (*88*).

We have used natural connected speech as auditory material. Although this adds to the ecological validity of our results, it makes it difficult to resolve the development of brain functions supporting multiple distinct aspects of language. For example, phrasal CTS taps in brain function supporting linguistic (syntactic, lexical, grammatical) as well as paralinguistic (prosody) information. Studies relying on carefully synthesized speech in which, e.g., prosody is removed (*27*).

Characterization of behavioral performance was suboptimal. We only asked simple comprehension questions, and comprehension scores suffered ceiling effects. This may explain why the link between CTS and behavior, which has been well documented in other studies (*18, 20, 22, 89*), was either weak (for phrasal nCTS) or non-significant (for syllabic nCTS) in our study despite having a sample size (*n* = 144) that largely surpasses that of previous studies. A more extensive neuropsychological assessment of language processing abilities could have further supported the behavioral relevance of the multiple developmental effects we identified.

## Conclusion

This study reveals distinct developmental trajectories for the neuronal processing of prosodic/syntactic (phrasal CTS) and syllabic/phonemic/acoustic information (syllabic CTS), and depending on the presence and the type of background noise. Overall, our results indicate that cortical processing of large linguistic units matures before that of smaller units, and that additional neuromaturational milestones need reaching for such processing to be optimal in adverse noise conditions. Unexpectedly, although visual speech information boosted the ability of the brain to track speech in noise, such boost was not affected by brain maturation. Finally, the ability to maintain phrasal tracking in noise was positively related to speech comprehension. These results therefore indicate that CTS tags behaviourally relevant neural mechanisms that progressively mature with age and experience following the trajectory presumed by grain-size proposals. Thus, the modulation of CTS by noise provides objective neurodevelopmental markers of multiple aspects of speech processing in noise.

## Material and Methods

### Participants

In total, 144 native French-speaking healthy right-handed children and young adults (age range: 5–27 years, 77 females) participated in this study. For some of the upcoming analyzes, participants were assigned to 5 age groups: 5–7 years (*n* = 31, 17 females), 7–8.5 years (*n* = 34, 17 females), 8.5–11.5 years (*n* = 28, 13 females), 11.5–18 years (*n* = 27, 15 females) and 18–27 years (*n* = 25, 15 females). Of note, the data collected from 73 of them was used in a previous study by our team (*32*). All participants had normal hearing according to pure-tone audiometry (i.e., hearing thresholds between 0 and 20 dB HL for 125, 250, 500, 1000, 2000, and 4000 and 8000 Hz), and normal dichotic perception, speech, and SiN perception for their age (data missing for the 20 youngest participants) according to another test assessing speech perception in noise (*79*).

The study had prior approval by the ULB-Hôpital Erasme Ethics Committee (Brussels, Belgium). Each participant or their legal representative gave written informed consent before participation. Participants were compensated with a gift card worth 25 euros for the neuroimaging assessment reported in the present study.

### Stimuli

The stimuli were derived from 12 audiovisual recordings of 4 native French-speaking narrators (2 females, 3 recordings per narrator) telling a story for ∼6-min (mean ± SD, 6.0 ± 0.8 min) Stories consisted of children’s fairy tales; for more details, see our previous report (*32*). In each video, the first 5 s were kept unaltered to enable participants to unambiguously identify the narrator’s voice and face they were requested to attend to. The remainder of the video was divided into 10 consecutive blocks of equal size that were assigned to 9 conditions. Two blocks were assigned to the *noiseless* condition in which the audio track was kept but the video was replaced by static pictures illustrating the story (mean ± SD picture presentation time across all videos, 27.7 ± 10.8 s). The remaining 8 blocks were assigned to 8 conditions in which the original sound was mixed with a background noise at 3 dB signal-to-noise ratio (SNR). This SNR was chosen as we assumed it was high enough to ensure children cope with the noise and keep their attention to the story, and low enough to introduce non-negligible interference; both assumptions proved accurate *a posteriori*. There were 4 different types of noise, and each type of noise was presented once with visual speech information (the original video), and once without visual speech information (static pictures illustrating the story). The different types of noise differed in the degree of energetic and informational interference they introduced (*90*). The least-energetic non-speech (i.e., non-informational) noise was a white noise high-pass filtered at 10000-Hz. The most-energetic non-speech noise had its spectral properties dynamically adapted to mirror those of the narrator’s voice ∼1 s around. The different-gender babble (i.e. informational) noise was a 5-talker cocktail party noise recorded by individuals of gender different from the narrator’s (i.e., a 5-male talker for female narrators, and vice-versa). The same-gender babble noise was a 5-talker cocktail party noise recorded by individuals of gender identical to the narrator’s. For both babble noises, the 5 individual noise components were obtained from a French audiobook database (http://www.litteratureaudio.com), normalized, and mixed linearly. The assignment of conditions to blocks was random, with the constraint that each of the 5 first and last blocks contained exactly 1 *noiseless* audio and each type of noise, 2 with visual speech and 2 without. Smooth audio and video transitions between blocks was ensured with 2-s fade-in and fade-out. Ensuing videos were grouped in 3 disjoint sets featuring one video of each of the narrators (total set duration: 23.0, 24.3, 24.65 min), and there were 4 versions of each set differing in condition random ordering.

### Experimental paradigm

During the imaging session, participants were laying on a bed with their head inside the MEG helmet. The lying position was chosen to maximize participants’ comfort and reduce head movements. Participants’ brain activity was recorded while they were attending 4 videos (separate recording for each video) of a randomly selected set and ordering of the videos presented in a random order, and finally while they were at *rest* (eyes opened, fixation cross) for 5 min. They were instructed to watch the videos attentively, listen to the narrators’ voice while ignoring the interfering noise, and remain as still as possible. After each video, they were asked 10 yes/no simple comprehension questions. Videos were projected onto a back-projection screen placed vertically, ∼120 cm away from the MEG helmet. The inner dimensions of the black frame were 35.2 cm (horizontal) and 28.8 cm (vertical), and narrators face spanned ∼15 cm (horizontal) and ∼20 cm (vertical). Participants could see the screen through a mirror placed above their head. In total the optical path from the screen to participants’ eyes was of ∼150 cm. Sounds were delivered at 60 dB (measured at ear-level) through a MEG-compatible front-facing flat-panel loudspeaker (Panphonics Oy, Espoo, Finland) placed ∼1 m behind the screen.

### Data acquisition

During the experimental conditions, participants’ brain activity was recorded with MEG at the CUB Hôpital Erasme. MEG was preferred to electroencephalography for its higher spatial resolution (*91*), and for its increased sensitivity to CTS (*4*). Neuromagnetic signals were recorded with a whole-scalp-covering MEG system (Triux, Elekta) placed in a lightweight magnetically shielded room (Maxshield, Elekta), the characteristics of which have been described elsewhere (*92*). The sensor array of the MEG system comprised 306 sensors arranged in 102 triplets of one magnetometer and two orthogonal planar gradiometers. Magnetometers measure the radial component of the magnetic field, while planar gradiometers measure its spatial derivative in the tangential directions. MEG signals were band-pass filtered at 0.1–330 Hz and sampled at 1000 Hz.

We used 4 head-position indicator coils to monitor subjects’ head position during the experimentation. Before the MEG session, we digitized the location of these coils and at least 300 head-surface points (on scalp, nose, and face) with respect to anatomical fiducials with an electromagnetic tracker (Fastrack, Polhemus).

Finally, subjects’ high-resolution 3D-T1 cerebral images were acquired with a magnetic resonance imaging (MRI) scanner (MRI 3T, Signa, General Electric) after the MEG session.

### Data preprocessing

Continuous MEG data were first preprocessed off-line using the temporal signal space separation method implemented in MaxFilter software (MaxFilter, Neuromag, Elekta; correlation limit 0.9, segment length 20 s) to suppress external interferences and to correct for head movements (*93, 94*). To further suppress physiological artifacts, 30 independent components were evaluated from the data band-pass filtered at 0.1–25 Hz and reduced to a rank of 30 with principal component analysis. Independent components corresponding to heartbeat, eye-blink, and eye-movement artifacts were identified, and corresponding MEG signals reconstructed by means of the mixing matrix were subtracted from the full-rank data. Across subjects and conditions, the number of subtracted components was 3.45 ± 1.23 (mean ± SD across subjects and recordings). Finally, time points at timings 1 s around remaining artifacts were set to bad. Data were considered contaminated by artifacts when MEG amplitude exceeded 5 pT in at least one magnetometer or 1 pT/cm in at least one gradiometer.

We extracted the temporal envelope of the attended speech (narrators’ voice) using a state-of-the-art approach (*95*). Briefly, audio signals were bandpass filtered using a gammatone filter bank (15 filters centered on logarithmically-spaced frequencies from 150 Hz to 4000 Hz), and subband envelopes were computed using Hilbert transform, elevated to the power 0.6, and averaged across bands.

### CTS quantification with accuracy of speech temporal envelope reconstruction

For each condition and participant, a global value of cortical tracking of the attended speech was evaluated for all left-hemisphere gradiometer sensors at once, and for all right-hemisphere gradiometer sensors at once. Using the mTRF toolbox (*54*), we trained a decoder on MEG data to reconstruct speech temporal envelope, and estimated its Pearson correlation with real speech temporal envelope. This correlation is often referred to as the reconstruction accuracy, and it provides a global measure of cortical tracking of speech (CTS).

The decoder tested on a given condition was built based on MEG data from all the other conditions. This procedure was preferred over a more conventional cross-validation approach in which the decoder is trained and tested on separate chunks of data from the same condition because of the paucity of data (*i.e.*, at most ∼2.4 min of data per condition). It is based on the rationale that the different conditions do modulate response amplitude but not its topography and temporal dynamics. In practice, electrophysiological data were band-pass filtered at 0.2–1.5 Hz (phrasal rate) or 2–8 Hz (syllabic rate), resampled to 10 Hz (phrasal) or 40 Hz (syllabic) and standardized. The decoder was built based on MEG data from –500 ms to 1000 ms (phrasal) or from 0 ms to 250 ms (syllabic) with respect to speech temporal envelope. Filtering and delay ranges were as in previous studies for phrasal (*4, 55*), and syllabic CTS (*10, 14, 31, 56*). Regularization was applied to limit the norm of the derivative of the reconstructed speech temporal envelope (*54*), by estimating the decoder for a fixed set of ridge values (λ = 2^-10^, 2^-8^, 2^-6^, 2^-4^, 2^-2^, 2^0^). The regularization parameter was determined with a classical 10-fold cross-validation approach: the data is split into 10 segments of equal length, the decoder is estimated for 9 segments and tested on the remaining segment, and this procedure is repeated 10 times until all segments have served as test segment. The ridge value yielding the maximum mean reconstruction accuracy is then retained. The ensuing decoder was then used to reconstruct speech temporal envelope in the left-out condition. Reconstruction accuracy was then estimated in 10 disjoint consecutive segments. We then retained the mean of this reconstruction accuracy, leaving us with one value for all combinations of subjects, conditions, hemispheres, and frequencies of interest.

### Normalized CTS in SiN conditions

Based on CTS values, we derived the normalized CTS (nCTS) in SiN conditions as the following contrast between CTS in SiN (*CTS*_SiN_) and noiseless (*CTS*_noiseless_) conditions:

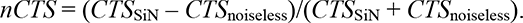

Such contrast presents the advantage of being specific to SiN processing abilities by factoring out the global level of CTS in the noiseless condition. However, it can be misleading when derived from negative CTS values (which may happen since CTS is an unsquared correlation value). For this reason, CTS values below a threshold of 10% of the mean CTS across all subjects, conditions and hemispheres were set to that threshold prior to nCTS computation. Thanks to this thresholding, the nCTS index takes values between –1 and 1, with negative values indicating that the noise reduces CTS.

### Developmental trajectory of CTS in noiseless and nCTS in SiN conditions

We used repeated measures ANOVA to assess the effect of brain hemisphere (left vs. right) and age group on CTS in noiseless conditions (dependent variable). This analysis was run separately for phrasal and syllabic CTS.

We used the same approach to analyze nCTS values in SiN conditions, this time with two additional factors: type of noise (least-energetic non-speech, most-energetic non-speech, different-gender babble vs. same-gender babble) and type of visual input (with vs. without visual speech). For both phrasal and syllabic nCTS, Mauchly sphericity tests indicated non-sphericity for the effect and interactions including the factor “type of noise” (*p* < 0.01). For this reason, Greenhouse-Geisser corrections were applied when needed.

For statistically significant effects involving age group, we used a model fitting approach to estimate the developmental trajectory of (n)CTS averaged across irrelevant factors. This approach is explained here for CTS, but the same was used for nCTS. We fitted to individual values of CTS three models involving different types of dependence on age:

Constant model: *CTS*(*age*) = *CTC*_constant_

Linear model: *CTS*(*age*) = *CTS*_0_ + *slope* × *age*

Logistic model: *CTS*(*age*) = *CTS*_min_ + (*CTS*_max_ – *CTS*_min_)/(1+exp(–*k*_a_ × (*age*–*age*_trans_))) The logistic model features an evolution of CTS with age from *CTS*_min_ to *CTS*_max_ with a transition at *age*_trans_ occurring at rate *k*_a_. Following this model, the maturation of CTS values roughly starts at *age*_trans_ – 2.2/*k*_a_ and finishes at *age*_trans_ + 2.2/*k*_a_, corresponding to 10 % and 90 % of the evolution from *CTS*_min_ to *CTS*_max_ (respectively). We also report on the percentage of increase in CTS, which is obtained as (*CTS*_max_ – *CTS*_min_)/*CTS*_min_ × 100 %.

Parameters were estimated with the least-square criterion, so that their values for the constant and linear models were trivial to obtain. Parameters of the logistic model were estimated with *fminsearch* Matlab function.

The models were compared statistically with a classical *F* test.

### Source reconstruction of CTS

As a preliminary step to estimate brain maps of CTS, MEG signals were projected into the source space. For that, MEG and MRI coordinate systems were co-registered using the 3 anatomical fiducial points for initial estimation and the head-surface points for further manual refinement. When a participant’s MRI was missing (*n* = 39), we used that of another participant of roughly the same age, which we linearly deformed to best match head-surface points using the CPD toolbox (*96*) embedded in FieldTrip toolbox (Donders Institute for Brain Cognition and Behaviour, Nijmegen, The Netherlands, RRID:SCR_004849) (*97*). The individual MRIs were segmented using Freesurfer software (Martinos Center for Biomedical Imaging, Boston, MA, RRID:SCR_001847) (*98*). Then, a non-linear transformation from individual MRIs to the MNI brain was computed using the spatial normalization algorithm implemented in Statistical Parametric Mapping (SPM8, Wellcome Department of Cognitive Neurology, London, UK, RRID:SCR_007037) (*99, 100*). This transformation was used to map a homogeneous 5-mm grid sampling the MNI brain volume onto individual brain volumes. For each subject and grid point, the MEG forward model corresponding to three orthogonal current dipoles was computed using the one-layer Boundary Element Method implemented in the MNE software suite (Martinos Centre for Biomedical Imaging, Boston, MA, RRID:SCR_005972) (*101*). The forward model was then reduced to its two first principal components. This procedure is justified by the insensitivity of MEG to currents radial to the skull, and hence, this dimension reduction leads to considering only the tangential sources. Source signals were then reconstructed with Minimum-Norm Estimates inverse solution (*102*).

We followed a similar approach to that used at the sensor level to estimate source-level CTS. For each grid point, we trained a decoder on the two-dimensional source time-series to reconstruct speech temporal envelope. Again, the decoder was trained on the data from all but one condition, and used to estimate CTS in the left-out condition. To speed up computation, the training was performed without cross-validation, with the ridge value retained in a sensor-space analysis run on all gradiometer sensors at once. This procedure yielded a source map of CTS for each participant, condition, and frequency range of interest; and because the source space was defined on the MNI brain, all CTS maps were inherently corregistered with the MNI brain. Hence, group-averaged maps were simply produced as the mean of individual maps within age groups, conditions and frequency ranges of interest.

We further identified the coordinates of local maxima in group-averaged CTS maps. Such local maxima of CTS are sets of contiguous voxels displaying higher CTS values than all neighbouring voxels. We only report statistically significant local maxima of CTS, disregarding the extents of these clusters. Indeed, cluster extent is hardly interpretable in view of the inherent smoothness of MEG source reconstruction (*103–105*).

Note that the adult MNI template was used in both children and adults despite the fact that spatial normalization may fail for brains of small size when using an adult template (*106*). However, this risk is overall negligible for the population studied here. Indeed, the brain volume does not change substantially from the age of 5 years to adulthood (*106*). This assumption has been confirmed by a study that specifically addressed this question in children aged above 6 years (*107*). This said, the precise anatomical location of anterior frontal and temporal opercular sources might be limited due to the greater deformation in those regions (*84*).

### Significance of local maxima of CTS

The statistical significance of the local maxima of CTS observed in group-averaged maps for each age group, condition and frequency range of interest was assessed with a non-parametric permutation test that intrinsically corrects for multiple spatial comparisons (*108*). First, participant and group-averaged *null* maps of CTS were computed with the MEG signals and the voice signal in each story rotated in time by about half of story length (i.e., the first and second halves were swapped, thereby destroying genuine coupling but preserving spectral properties). The exact temporal rotation applied was chosen to match a pause in speech to enforce continuity. Group-averaged difference maps were obtained by subtracting *genuine* and *null* group-averaged CTS maps. Under the null hypothesis that CTS maps are the same whatever the experimental condition, the labeling *genuine* or *null* are exchangeable prior to difference map computation (*108*). To reject this hypothesis and to compute a significance level for the correctly labeled difference map, the sample distribution of the maximum of the difference map’s absolute value within the entire brain was computed from a subset of 1000 permutations. The threshold at *p* < 0.05 was computed as the 95 percentile of the sample distribution (*108*). All supra-threshold local maxima of CTS were interpreted as indicative of brain regions showing statistically significant CTS and will be referred to as sources of CTS.

Permutation tests can be too conservative for voxels other than the one with the maximum observed statistic (*108*). For example, dominant CTS values in the right auditory cortex could bias the permutation distribution and overshadow weaker CTS values in the left auditory cortex, even if these were highly consistent across subjects. Therefore, the permutation test described above was conducted separately for left- and right-hemisphere voxels.

### Effect of age group and conditions on CTS source location

We evaluated for each frequency range if sources of CTS tended to cluster according to some categories. Five different categories were considered: (i) age-group category (5 age groups), (ii) visual category (with vs. without visual input), (iii) 3-noise category (noiseless vs. non-speech noises vs. babble noises), (iv) 2-noise category (noiseless and non-speech noises vs. babble noises), and (v) presence of noise category (noiseless vs SiN). For this analysis, we gathered the coordinates of all sources of CTS in all conditions (8 SiN and 2 instances of noiseless speech). For each (target) source and category we computed the proportion of the 10 closest sources (excluding those for the same condition within the same age group as the target source) sharing the same category as the target source, we divided that proportion by that expected by chance (i.e., the total number of sources sharing the same category as the target source divided by the total number of sources), subtracted 1, and multiplied by 100 %. The mean of these values for a given category across all sources indicates the increase in chance (in percent; compared with what is expected by chance) of finding another CTS source of that category in the close vicinity. For statistical assessment, this mean value was compared with its permutation distribution where the CTS sources were assigned to random labels (1000 permutations).

## Acknowledgments

**Funding:** Florian Destoky, Julie Bertels and Mathieu Bourguignon have been supported by the program Attract of Innoviris (grants 2015-BB2B-10 and 2019-BFB-110). Julie Bertels has been supported by a research grant from the Fonds de Soutien Marguerite-Marie Delacroix (Brussels, Belgium). Maxime Niesen has been supported by the Fonds Erasme (Brussels, Belgium). Xavier De Tiège is Post-doctorate Clinical Master Specialist at the Fonds de la Recherche Scientifique (F.R.S.-FNRS, Brussels, Belgium). Mathieu Bourguignon has been supported by the Marie Skłodowska-Curie Action of the European Commission (grant 743562). The MEG project at the CUB Hôpital Erasme and this study were financially supported by the Fonds Erasme (Research convention “Les Voies du Savoir”, Brussels, Belgium). The PET-MR project at the CUB Hôpital Erasme is supported by the Association Vinçotte Nuclear (AVN, Brussels, Belgium).

## Author contributions

Conceptualization: JB, FD, TC, MVG, MBa, NM, XDT, MBo

Methodology: JB, MN, VW, MBo

Investigation: JB, MN, FD, AR, NT, MBo

Supervision: XDT, MBo

Writing—original draft: JB, MN, TC

Writing—review & editing: FD, MVG, VW, AR, NT, MBa, NM, XDT, MBo

## Data and materials availability

All data, code, and materials used in the analyses are available at the following link: https://osf.io/4uzrm/

## Supplementary Material

**Table S1.** Results of the ANOVAs run on CTS in the noiseless condition.

|  | Phrasal CTS |  |  |  | Syllabic CTS |  |  |  |
| --- | --- | --- | --- | --- | --- | --- | --- | --- |
|  | *F* | df1 | df2 | *p* | *F* | df1 | df2 | *p* |
| Hemisphere | **33.08** | **1** | **139** | **<0.0001** | **31.83** | **1** | **139** | **<0.0001** |
| Age | 0.94 | 4 | 139 | 0.44 | 1.76 | 4 | 139 | 0.14 |
| Age x Hemisphere | 0.70 | 4 | 139 | 0.59 | **2.91** | **4** | **139** | **0.0238** |

**Table S2.** Results of the ANOVAs run on nCTS in noisy conditions.

|  | Phrasal CTS |  |  |  | Syllabic CTS |  |  |  |
| --- | --- | --- | --- | --- | --- | --- | --- | --- |
|  | *F* | df1 | df2 | *p* | *F* | df1 | df2 | *p* |
| Noise | **218** | **1.73** | **241** | **< 0.0001** | **119** | **2.77** | **384** | **< 0.0001** |
| Hemisphere | **13.1** | **1** | **139** | **0.0004** | **3.69** | **1** | **139** | **0.057** |
| Visual input | **160** | **1** | **139** | **< 0.0001** | **31.38** | **1** | **139** | **< 0.0001** |
| Age | **13.3** | **4** | **139** | **< 0.0001** | 1.25 | 4 | 139 | 0.29 |
| Hemisphere x Noise | **9.01** | **2.10** | **292** | **< 0.0001** | 0.68 | 2.80 | 389 | 0.55 |
| Visual input x Noise | **29.7** | **2.61** | **263** | **< 0.0001** | 0.99 | 2.82 | 392 | 0.40 |
| Age x Noise | **11.9** | **6.92** | **241** | **< 0.0001** | 1.36 | 11.1 | 384 | 0.19 |
| Age x Hemisphere x Noise | 0.95 | 8.40 | 292 | 0.48 | **1.87** | **11.2** | **389** | **0.041** |
| Age x Visual input | 0.90 | 4 | 139 | 0.47 | 0.95 | 4 | 139 | 0.44 |
| Age x Visual input x Noise | 1.21 | 10.4 | 363 | 0.28 | 0.72 | 11.3 | 392 | 0.72 |

**Table S3.** Parametric models of the dependence on age of phrasal nCTS (averaged across hemispheres) and syllabic nCTS (averaged across least- and most-energetic conditions or across different- and same-gender conditions) in conditions with and without visual speech.

| Noise type | Constant vs. Linear |  | Linear vs. Logistic |  | Constant vs. Logistic |  | Model |
| --- | --- | --- | --- | --- | --- | --- | --- |
|  | *F*(1,142) | *p* | *F*(2,140) | *p* | *F*(3,140) | *p* | *nCTS*(age) = |
| Phrasal nCTS without visual speech |  |  |  |  |  |  |  |
| least-energetic non-speech | 1.62 | 0.21 | 0.17 | 0.85 | 0.64 | 0.59 | -0.003 |
| most-energetic non-speech | 2.43 | 0.12 | 1.19 | 0.31 | 1.61 | 0.19 | -0.009 |
| different-gender babble | 20.9 | <0.0001 | 8.76 | 0.0003 | 13.57 | <0.0001 | $-0.39 + 0.25 / (1 + \exp(-1.4(\text{age} - 7.3)))$ |
| same-gender babble | 34.1 | <0.0001 | 9.12 | 0.0002 | 18.74 | <0.0001 | $-0.43 + 0.28 / (1 + \exp(-1.4(\text{age} - 7.9)))$ |
| Phrasal nCTS with visual speech |  |  |  |  |  |  |  |
| least-energetic non-speech | 1.60 | 0.21 | 1.07 | 0.35 | 1.25 | 0.29 | -0.005 |
| most-energetic non-speech | 4.37 | 0.038 | 6.52 | 0.0020 | 5.92 | 0.0008 | $-0.03 + 0.06 / (1 + \exp(-1072(\text{age} - 6.8)))$ |
| different-gender babble | 23.6 | <0.0001 | 9.47 | 0.0001 | 15.1 | <0.0001 | $-0.39 + 0.34 / (1 + \exp(-0.89(\text{age} - 6.7)))$ |
| same-gender babble | 22.9 | <0.0001 | 11.6 | <0.0001 | 16.5 | <0.0001 | $-0.33 + 0.27 / (1 + \exp(-1.1(\text{age} - 7.3)))$ |
| Syllabic nCTS without visual speech |  |  |  |  |  |  |  |
| non-speech, left hem | 6.35 | 0.013 | 1.22 | 0.30 | 2.94 | 0.035 | $-0.080 + 0.0054 \times \text{age}$ |
| babble, left hem | 3.00 | 0.086 | 2.29 | 0.11 | 2.54 | 0.059 | $-0.21 \quad [-0.27 + 0.0050 \times \text{age}]$ |
| non-speech, right hem | 0.79 | 0.38 | 0.45 | 0.64 | 0.56 | 0.64 | 0.0035 |
| babble, right hem | 3.66 | 0.058 | 0.12 | 0.89 | 1.28 | 0.28 | $-0.18 \quad [-0.24 + 0.0052 \times \text{age}]$ |
| Syllabic nCTS with visual speech |  |  |  |  |  |  |  |
| non-speech, left hem | 1.94 | 0.17 | 0.00 | 1.00 | 0.64 | 0.59 | $0.025 \quad [-0.009 + 0.0029 \times \text{age}]$ |
| babble, left hem | 8.51 | 0.0041 | 0.20 | 0.82 | 2.94 | 0.036 | $-0.24 + 0.0080 \times \text{age}$ |
| non-speech, right hem | 2.28 | 0.13 | 0.00 | 1.00 | 0.75 | 0.53 | 0.042 |
| babble, right hem | 2.89 | 0.091 | 0.40 | 0.67 | 1.22 | 0.30 | $-0.12 \quad [-0.17 + 0.0045 \times \text{age}]$ |

**Fig. S1.**
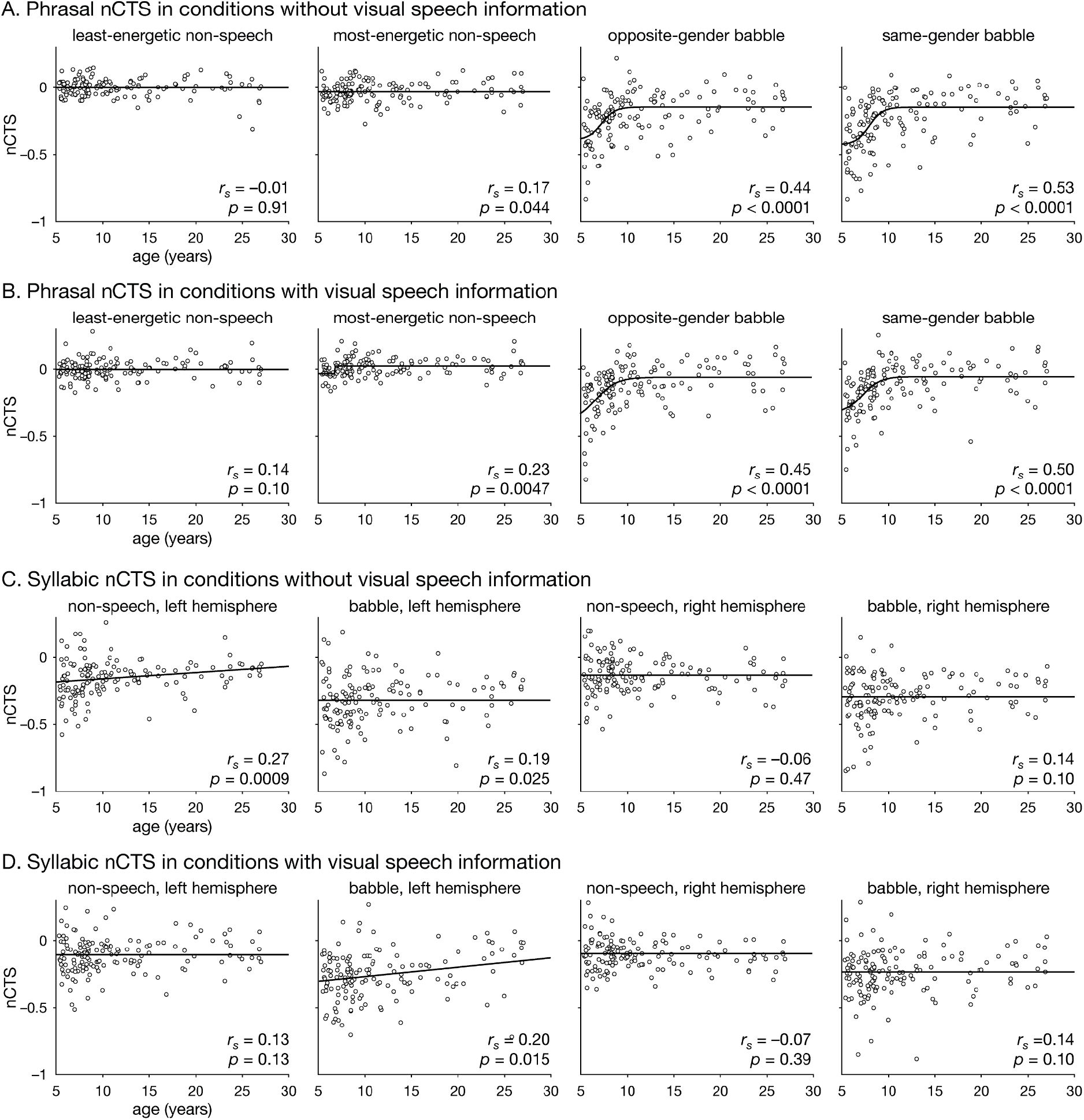
Dependence on age of phrasal nCTS (averaged across hemispheres) in conditions without (A) and with visual speech (B), and of syllabic nCTS (averaged across least- and most-energetic conditions or across different- and same-gender conditions) in conditions without (C) and with visual speech (D).

**Fig. S2.**
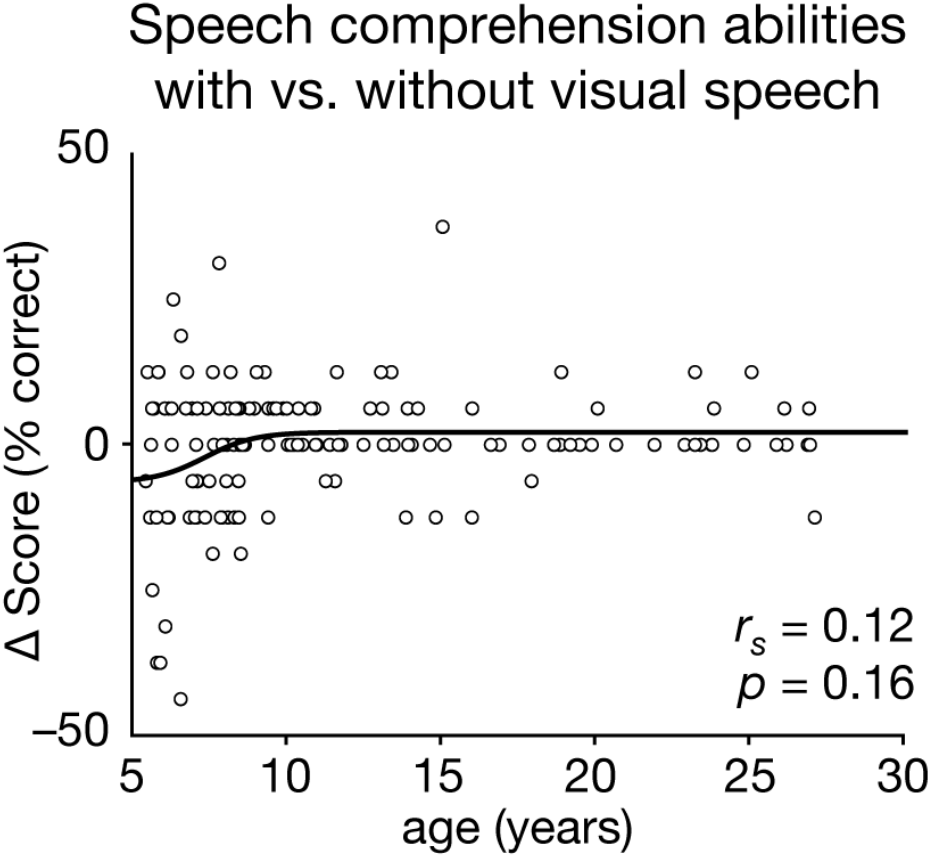
Dependence on age of speech comprehension scores contrasted between conditions with vs. without visual speech information. Although Spearman correlation was not statistically significant, a non-linear model explained significantly more variance than a constant model (*F*(3,138) = 2.94, *p* = 0.035), and marginally more than a linear model (*F*(2,138) = 2.84, *p* = 0.062), which itself explained marginally more variance than a constant model (*F*(1,138) = 3.07, *p* = 0.082). According to the non-linear model (score(age) = –0.065 + 0.086/(1+exp(–1.2(age–7.3)))), the gain in comprehension afforded by visual speech increased as a function of age, and was positive only from age 8.2.

## S1 Discussion. Beneficial effect of visual speech on CTS in noise

The beneficial effect of synchronized visual speech for speech perception and comprehension is largely documented, especially in noise conditions (*41, 109*). It was also observed on CTS in adults (*34, 37, 110, 111*). In our study, visual speech boosted phrasal CTS mainly in babble noise conditions (but also in the most-energetic non-speech condition) and syllabic CTS in all noise conditions with no effect of age. This suggests that the brain would leverage visual speech to parse speech into syllables no matter what the listening conditions are, while it would use such visual information to parse speech into phrases only when parsing is made difficult by challenging competing noise. This distinction nicely echoes the view that audiovisual integration dissociates into two modes, one in which vision and audition provide complementary information and one in which they provide redundant information (*112*). Given that parsing a continuous speech stream into meaningful units is a difficult task, the brain would always strive to combine the information from both modalities. The complementary mode of audiovisual speech integration would be based on the extraction of relevant features of the mouth configuration to derive phonetic information, complementing those derived from acoustic speech signals (*112*). Parsing of phrases based on auditory information is typically easier because listeners can rely on three main prosodic cues: pitch change, final lengthening, and pauses (*65*). Accordingly, visual speech information about phrase boundaries, which is at best redundant with auditory speech information, would be of use for phrasal parsing only when access to these cues is compromised, as in challenging noise conditions. This complementary mode of audiovisual speech integration would rather be based on the extraction of the temporal dynamics of lip, jaw and head movements to support speech parsing (*112*).

## S2 Discussion. Impact of noise properties on the neural network for CTS

Overall, sources of phrasal CTS were more widely distributed in language-related areas (*113*) than those of syllabic CTS. This differential recruitment may reflect the increased reliance on top-down information from higher-order language areas to facilitate speech processing at the phrasal level compared to syllabic level (*114*).

The dominant sources of syllabic CTS clustered around Heschel’s gyrus bilaterally, while those of phrasal CTS located in middle and posterior STG, in line with previous reports (*17, 104*). Interestingly, STG sources of phrasal CTS extended more anteriorly in the informational noise condition compared with the other conditions. This is in line with the existence of a posterior-to-anterior gradient in the STG with increasing complexity of the auditory stimulus (*115*).

Sources of phrasal CTS also located in ventral posterior temporal areas, attributed to semantic aspects (*113*) and sentence-level processing (*116*). However, this recruitment of the ventral stream was essentially restricted to the informational noise condition, probably reflecting the reliance on semantic processes to correctly parse phrasal boundaries in challenging conditions.

In the frontal lobe, the reliance on semantic processing in phrasal CTS may be reflected by the anterior shift of sources towards IFG pars orbitalis in informational noise, consistent with the functional segmentation of the IFG characterized by a posterior-dorsal (phonology) to anterior-ventral (semantics) gradient (*117–119*). However, the presence of syllabic CTS sources in the anterior-ventral IFG is somewhat at odds with this functional segregation of the IFG. Tentatively, it might relate to lower-level semantic processes, such as those potentially needed to predict plausible syllabic sequences. Finally, the dorsal part of the pars opercularis was reported to be involved in syntactic processing (*116*). Our data fit well into this perspective, since phrasal but not syllabic CTS sources localized in this region, and only in the least-challenging conditions. In more challenging conditions, emphasis is expected to be placed on semantic rather than syntax, as exemplified by suppressed evoked responses to syntactic violations in challenging noisy conditions (*120*).

